# Genetic lineage of the Amami islanders inferred from classical genetic markers

**DOI:** 10.1101/2021.04.18.440379

**Authors:** Yuri Nishikawa, Takafumi Ishida

**Author notes:** Correspondence: Yuri Nishikawa, Department of Biological Sciences, Graduate School of Science, The University of Tokyo, Hongo 7-3-1, Bunkyo-ku, Tokyo 113-0033, Japan.

## Abstract

The genetic structure of the people of mainland Japan and Okinawa has been gradually unveiled in recent years. However, previous anthropological studies dealing with people in the Amami islands, located between mainland Japan and Okinawa, were less informative because of the lack of genetic data. In this study, we collected DNAs from 104 subjects in two of the Amami islands, Amami-Oshima island and Kikai island. We analyzed the D-loop region of mtDNA, four Y-STRs, and four autosomal nonsynonymous SNPs to clarify the Amami islanders’ genetic structure compared with peoples in Okinawa, mainland Japan, and other regions of East Asia. We found that the Amami islanders showed a genetically intermediate position between mainland Japan and Okinawa in mtDNA and Y-STR. However, the frequencies of several autosomal SNPs in the Amami islanders indicated a significant difference from mainland Japanese, which may be because of the gene flow from Okinawa but not natural selection. Moreover, extremely high or low frequencies of several alleles implied a founder effect in Kikai islanders. Note that there is room for the interpretation of the results because of the small sample size and number of alleles in the present study. Geographically broad and detailed samplings and genome-wide analyses are awaited.

## 1. Introduction

The genetic structure of the people of Japan has been gradually unveiled in recent years. The dual structure model (Hanihara, 1991) is a well-known hypothesis that predicts that the modern Japanese population was formed by the admixture of two populations, Jomon and Yayoi; Jomon peoples dwelled in the Japanese Archipelago since about 18,000 years ago, and Yayoi peoples immigrated from the Asian continent 2,000–3,000 years ago (Habu, 2004; Nakagome et al., 2015). This model is also supported largely by genomic/genetic studies (Jinam et al., 2012; Nakagome et al., 2015). According to this model, there are two populations assumed to be strongly contributed from Jomon peoples. One of these populations is the Ainus, indigenous in Hokkaido, the most northern part of Japan. The other population comprises people living in the Ryukyu Archipelago, southern end of Japan (Fig. 1).

**Fig. 1.**
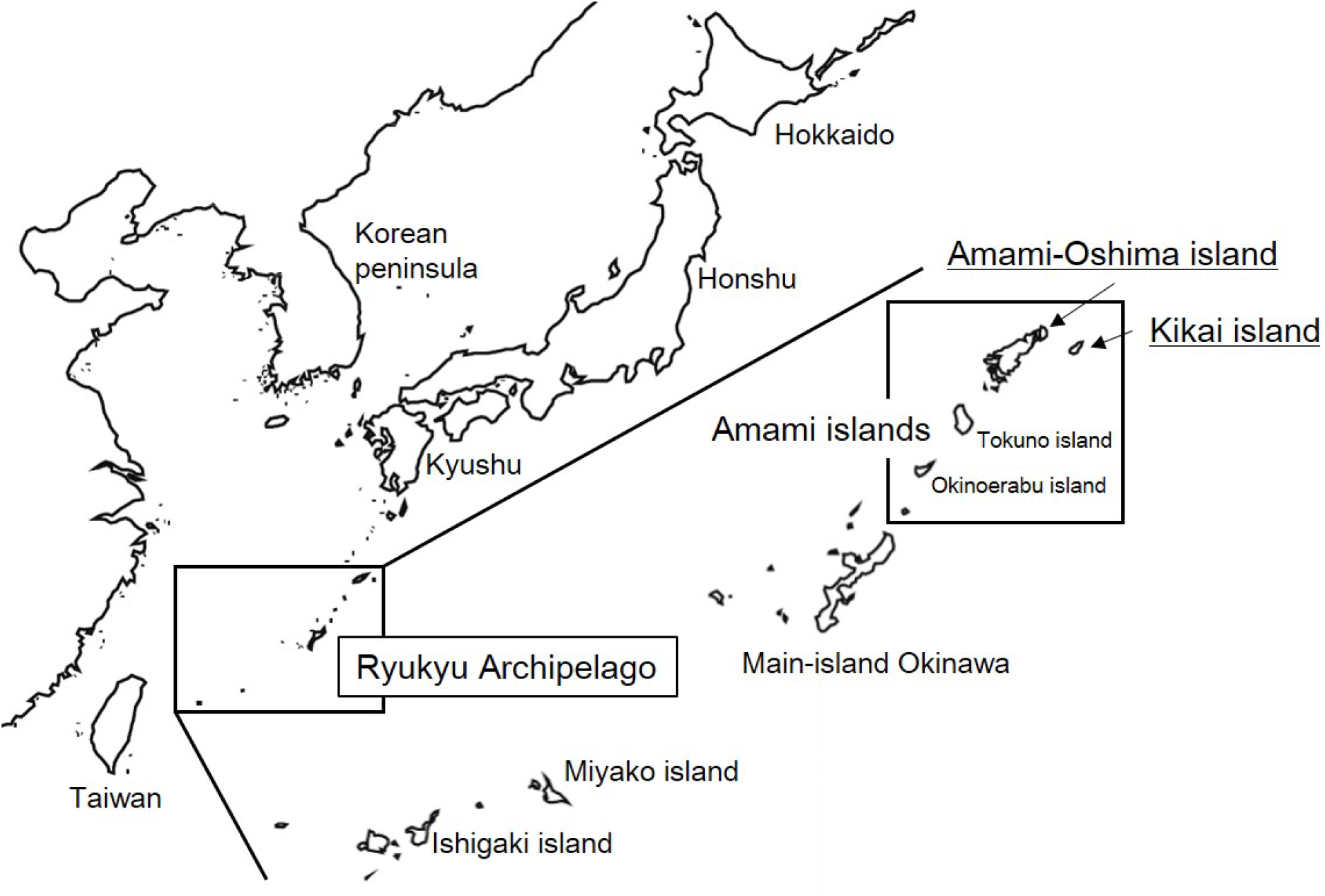
Map of East Asia. Amami-Oshima island and Kikai island are placed on the northern edge of the Ryukyu Archipelago. Main-island Okinawa, Miyako island and Ishigaki island are included in Okinawa. Hokkaido, Honshu and Kyushu are included in mainland Japan (Hondo).

This model has been supported by previous genetic studies among the Ryukyuans, Ainus, mainland Japanese, and other East Asian populations. Genetic similarity between the Ryukyuans and Ainus has been observed (Hammer et al., 2006; Koganebuchi et al. 2012), whereas the Ryukyuans are believed to be differentiated from mainland Japanese by the analyses of the Y chromosome (Hammer & Horai, 1995; Hammer et al., 2006), mtDNA (Horai et al., 1996; Tanaka et al., 2004), autosomal markers (Omoto & Saitou, 1997), and genome-wide SNP data (Yamaguchi-Kabata et al., 2008). Additionally, modern Ryukyuans are believed to have no direct genetic relations with the Taiwan aborigines (Matsukusa et al., 2010) or human skeletons in the late Pleistocene (Sato et al., 2014; Mizuno et al., 2021) that are older than Jomon peoples found in several sites in the Ryukyu Archipelago (Kobayashi et al., 1974; Nakagawa et al., 2010). Furthermore, rich genetic variations among the residents on the islands of the Ryukyu Archipelago have been observed (Sato et al., 2014; Matsunami et al., 2021).

However, in those studies mentioned above, much attention was not paid to the Amami islands located in the northern Ryukyu Archipelago (Fig. 1). There have been historical differences between the Amami islands and the rest of the Ryukyu Archipelago (Okinawa prefecture). Some researchers claim that the Amami islands played important roles in the trade among East Asia, including mainland Japan and China, around the 7th–11th century, which distinguished the Amami islands from Okinawa (Takanashi, 2011; Yoshinari, 2011). In Kikai island, one of the Amami islands, there is the Gusuku Site group which is supposed to be the local agency of the government of mainland Japan in the 9th–13th century (Yoshinari, 2011; Takamiya et al., 2019), and this corresponds to the hypothesis that there was population migration into the Ryukyu Archipelago from the north via the Amami islands around the 11th–12th century (Takamiya, 2013). In the 15th century, the Amami islands were conquered by the Ryukyu Kingdom, which was an independent country centered on main-island Okinawa (Yoshinari & Fuku, 2007), but since 1609, the Amami islands were directly ruled by the Satsuma-Han, one of the feudal domains in mainland Japan; this has brought about administrative and cultural differences between the Amami islands and Okinawa (Tsuha, 2012).

Despite these non-negligible historical and cultural features, few anthropological studies, including genetic studies about the Amami islanders, have been conducted. Dodo et al. (1998) analyzed 22 cranial nonmetric traits of early modern Amami and Okinawa islanders and found that they shared morphological similarities. Nishiyama et al. (2012) indicated a low but significant level of genetic difference between the Amami islanders and mainland Japanese using SNP data, but their research was limited to only two islands, Tokuno and Okinoerabu islands (Fig. 1), in the southern part of the Amami islands, and they concluded that studies that cover further north areas should be indispensable.

We then newly focused on two of the Amami islands, Amami-Oshima and Kikai islands, in this research. Amami-Oshima island (Fig. 1) is the largest island and has the largest population in the Amami islands (Kagoshima Prefecture, 2018a). Kikai island (Fig. 1) is a relatively small island with a population of about 7,000 (Kagoshima Prefecture, 2018b), located on the east of Amami-Oshima island, and assumed to have played a very important role in the relationship between the Ryukyu Archipelago and mainland Japan around the 11th–12th century (Takanashi, 2009; Yoshinari, 2011; Takamiya, 2013; Takamiya et al., 2019). On the basis of the previous studies and historical backgrounds described above, we hypothesized that people in these two islands might genetically be located in the intermediate position between Okinawa and mainland Japan. This study aims to clarify the genetic structure of the Amami islanders compared with peoples in Okinawa, mainland Japan, and other regions in East Asia.

## 2. Materials and methods

### 2.1. Subjects

A total of 104 Amami islanders (Fig. 1), 78 in Amami-Oshima island (49 males and 29 females) and 26 in Kikai island (15 males and 11 females), who were not genetically related, were the subjects of this study. This research was approved by the Research Ethics Committee of the University of Tokyo, and all subjects provided informed consent. We collected nail or oral mucosa samples from the subjects. Their origins were confirmed by an interview that at least one of the grandparents of each subject was born in the Amami islands.

### 2.2. DNA extraction

We extracted DNA from nail samples using the ISOHAIR (Nippon Gene) and oral mucosa samples using the High Pure PCR Template Preparation Kit (Roche).

### 2.3. Genotyping

We amplified the D-loop region (16024–16569 and 1–41 of the rCRS; Anderson et al., 1981; Andrews et al., 1999) of mtDNA by nested PCR to avoid nuclear insert contaminations. The first PCR for 9 kb of mtDNA including the D-loop region was conducted (primers: GENBANK_MT_26F & GENBANK_MT_5R) with an initial denaturation at 94°C for 2 min, followed by 25 cycles of denaturation at 98°C for 10 s, annealing at 72°C for 30 s, extension at 68°C for 10 min. The second PCR for the D-loop region was conducted (primers: L15996F & H408R) with an initial denaturation at 94°C for 2 min, followed by 25 cycles of denaturation at 98°C for 10 s, annealing at 56°C for 30 s, extension at 68°C for 1 min. After confirmation by 2% agarose gel electrophoresis, PCR products were purified using the PCR Clean-Up Mini Kit (FAVORGEN) and sequenced by outsourcing those to Fasmac (Kanagawa, Japan) or Eurofins Genomics (Tokyo, Japan). The primers used for PCR amplification and sequencing are listed in Supp. Table 1.

Four Y-STRs (DYS393, DYS19, DYS391, and DYS438) of which genetic diversities differed between Honshu (Fig. 1) and Okinawa (Uchihi et al., 2003) were selected. PCRs for these loci were conducted with an initial denaturation at 95°C for 10 min, followed by 45 cycles of denaturation at 95°C for 15 s, annealing at 53°C–59°C for 30 s, extension at 72°C for 1 min, followed by a final extension of 5 min at 72°C. After confirmation by 2% agarose gel electrophoresis, PCR products were purified using the PCR Clean-Up Mini Kit (FAVORGEN) and sequenced by outsourcing those to Fasmac (Kanagawa, Japan) or Eurofins Genomics (Tokyo, Japan). The primers and annealing temperatures for each PCR amplification are listed in Supp. Table 2.

We genotyped four nonsynonymous autosomal SNPs (rs3827760, rs17822931, rs2070235, and rs14103), of which genotype frequencies differed between the Hondo cluster, which includes most of the individuals in mainland Japan and the Ryukyu cluster, which includes most of the individuals in Okinawa (Yamaguchi-Kabata et al., 2008) by PCR-RFLP. PCRs were conducted with an initial denaturation at 95°C for 10 min, followed by 40 cycles of denaturation at 95°C for 15 s, annealing at 54°C–65°C for 30 s, extension at 72°C for 1 min, followed by a final extension of 5 min at 72°C. After confirmation by 2% agarose gel electrophoresis, PCR products were purified using the PCR Clean-Up Mini Kit (FAVORGEN) and digested with an appropriate restriction enzyme at 37°C for 1 h. Thereafter, we electrophoresed these digested products on 4% agarose gel for genotype determination. The primers, annealing temperatures, and restriction enzymes for each locus are listed in Supp. Table 3.

### 2.4. Data analyses

The mtDNA sequences were aligned using MEGA X (Kumar et al., 2018) together with sequences of other Asian populations downloaded from the NCBI (National Center for Biotechnology Information) database. Net genetic distances (*d*_*A*_) between populations were calculated from these sequences under the Tamura-Nei model (Tamura & Nei, 1993) with gamma distribution. Phylogenetic networks between populations based on *d*_*A*_ distances were constructed in the Neighbor-Net method (Bryant & Moulton, 2004) by using SplitsTree4 (Huson & Bryant, 2006). Pairwise *F*_*ST*_ between regions and nucleotide diversities within each population were calculated by using Arlequin ver 3.5.2.2 (Excoffier & Lischer, 2010), and Tajima’s *D* (Tajima, 1989) within each population were calculated using DnaSP 5.10 (Librado & Rozas, 2009). The probabilities of Tajima’s *D* were computed through 5,000 coalescence simulations.

We aligned the Y-STR sequences using MEGA X and counted the number of repeats. Pairwise *R*_*ST*_ between regions was calculated by using Arlequin ver 3.5.2.2 as well as gene diversities.

Fisher’s exact test was used to compare the allele frequencies of the SNPs between populations.

## 3. Results

### 3.1. mtDNA

D-loop sequences (587 bp) were determined for a total of 102 subjects whose maternal grandmothers had been born in the Amami islands. Fig. 2 and Supp. Fig. 1 show the phylogenetic networks for 78 Amami-Oshima islanders, 24 Kikai islanders, and other Asian populations. Amami-Oshima and Kikai islands were close to each other in both networks, and the Amami islanders did not form a clear cluster with the residents of Honshu, Kyushu, the Okinawa islands, or any other populations. Table 1 shows the nucleotide diversities and Tajima’s *D* in each population. Nucleotide diversities in the Amami islands were almost the same as those in other regions in Japan. Tajima’s *D* values were significantly negative in the Amami islands (−1.613, *P* = 0.024) and Amami-Oshima island (−1.539, *P* = 0.038). Tajima’s *D* value in Kikai island was negative but not significant (−1.494, *P* = 0.050). Table 2 shows the pairwise *F*_*ST*_ values between regions. *F*_*ST*_ value between Amami-Oshima and Kikai islands was not significant. *F*_*ST*_ values between Amami-Oshima island and Honshu, between Amami-Oshima island and Okinawa, between Amami-Oshima and Ishigaki islands, and between Kikai and Ishigaki islands were significant. Additionally, *F*_*ST*_ values between Korea and four regions in Japan (Amami-Oshima island, Kikai island, Kyushu, and Honshu) were not significant.

**Fig. 2.**
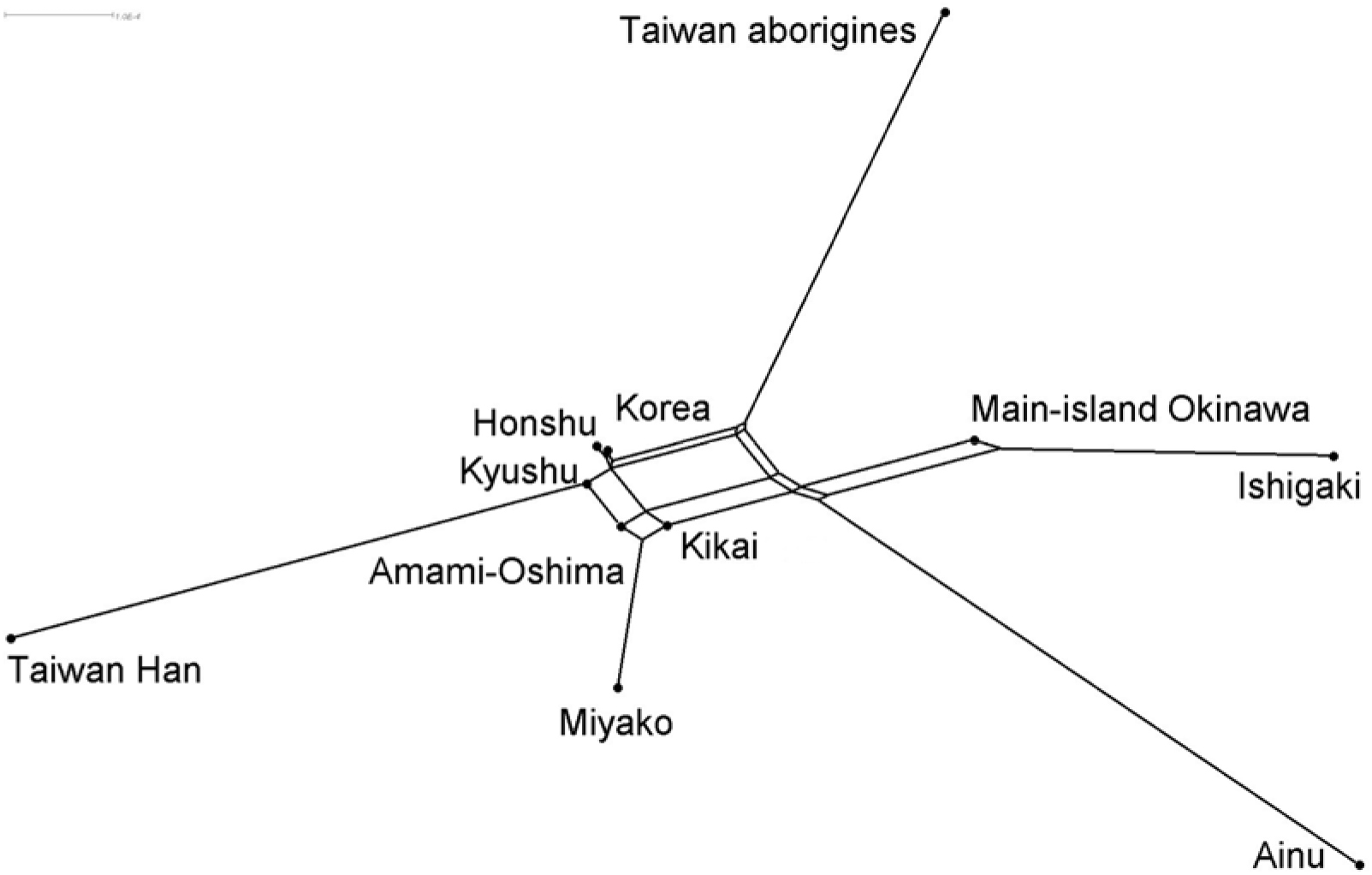
Phylogenetic network between populations based on *d*_*A*_ distances calculated from 337 bp of D-loop region. Data of Miyako, Ishigaki and Main-island Okinawa were taken from Matsukusa et al. (2010). Data of Honshu were taken from Oota et al. (2002). Data of Ainu, Korea and Taiwan Han were taken from Horai et al. (1996). Data of Taiwan aborigines were taken from Tajima et al. (2003).

**Table 1.** Nucleotide diversities and Tajima’s *D* values of mtDNA

| | Size | Nucleotide diversity | Tajima's $D$ | $P$value | Ref. |
| --- | --- | --- | --- | --- | --- |
| Amami islands | 102 | $0.019 \pm 0.010$ | -1.613 | 0.024 | Present study |
| Amami-Oshima island | 78 | $0.019 \pm 0.010$ | -1.516 | 0.042 | Present study |
| Kikai island | 24 | $0.020 \pm 0.011$ | -1.494 | 0.050 | Present study |
| Main island Okinawa | 95 | $0.019 \pm 0.010$ | -1.972 | 0.004 | Matsukusa et al., 2010 |
| Miyako island | 66 | $0.015 \pm 0.008$ | -1.916 | 0.008 | Matsukusa et al., 2010 |
| Ishigaki island | 63 | $0.016 \pm 0.008$ | -1.582 | 0.039 | Matsukusa et al., 2010 |
| Honshu | 89 | $0.015 \pm 0.008$ | -2.050 | 0.002 | Oota et al., 2002 |

For the analyses of mtDNA, one subject from Kikai island whose maternal grandmother was born in Amami-Oshima island was excluded from Kikai island samples and included in Amami-Oshima island samples.

**Table 2.** Pairwise *F*_*ST*_ values of mtDNA

|  | Amami-Oshima island | Kikai island | Main-island Okinawa | Miyako island | Ishigaki island | Kyushu | Honshu | Ainu | Korea | Taiwan Han | Taiwan aborigines |
| --- | --- | --- | --- | --- | --- | --- | --- | --- | --- | --- | --- |
| Amami-Oshima island (n=78) | 0 |  |  |  |  |  |  |  |  |  |  |
| Kikai island (n=24) | -0.007 | 0 |  |  |  |  |  |  |  |  |  |
| Main-island Okinawa (n=95) | **0.026** | 0.003 | 0 |  |  |  |  |  |  |  |  |
| Miyako island (n=66) | 0.004 | -0.001 | **0.034** | 0 |  |  |  |  |  |  |  |
| Ishigaki island (n=63) | **0.044** | **0.028** | **0.013** | **0.043** | 0 |  |  |  |  |  |  |
| Kyushu (n=104) | -0.001 | -0.004 | **0.036** | **0.013** | **0.055** | 0 |  |  |  |  |  |
| Honshu (n=89) | **0.009** | 0.010 | **0.043** | **0.012** | **0.045** | **0.008** | 0 |  |  |  |  |
| Ainu (n=51) | **0.043** | **0.027** | **0.043** | **0.062** | **0.068** | **0.042** | **0.074** | 0 |  |  |  |
| Korea (n=64) | 0.004 | -0.002 | **0.036** | **0.014** | **0.048** | -0.001 | 0.004 | **0.040** | 0 |  |  |
| Taiwan Han (n=52) | **0.033** | **0.037** | **0.081** | **0.053** | **0.102** | **0.030** | **0.033** | **0.078** | **0.039** | 0 |  |
| Taiwan aborigines (n=180) | **0.074** | **0.057** | **0.097** | **0.103** | **0.120** | **0.060** | **0.092** | **0.080** | **0.069** | **0.084** | 0 |

### 3.2. Y-STR

Y-STR sequences were determined for 58 male subjects whose paternal grandfathers had been born in the Amami islands. Twenty-five different haplotypes were observed in the Amami islands (Supp. Table 4). Three of them were observed in Kikai island but not in Amami-Oshima island. The paternal grandfather of one subject who had one of the three haplotypes (Ht11) was born in Amami-Oshima island. This subject was counted as Amami-Oshima samples in the Y-STR analyses. Comparing with other regions in Japan (Hashiyada et al., 2008), six haplotypes observed in Amami-Oshima island were not in Okinawa, and one haplotype was found only in Amami-Oshima island and Okinawa but not in any other regions. The allele frequency distributions of four loci were similar among regions (Supp. Table 5). Gene diversities of the four loci are shown in Supp. Table 6. In Amami-Oshima island, gene diversities of any loci did not significantly differ with Honshu or Okinawa. In Kikai island, gene diversities of the loci except DYS438 did not differ with other regions, and that of DYS438 was significantly higher than any other regions. Table 3 shows the pairwise *R*_*ST*_ values between regions. *R*_*ST*_ value between Amami-Oshima and Kikai islands was significant. *R*_*ST*_ values between Amami-Oshima island and Okinawa and between Kikai island and Okinawa were not significant.

**Table 3.**
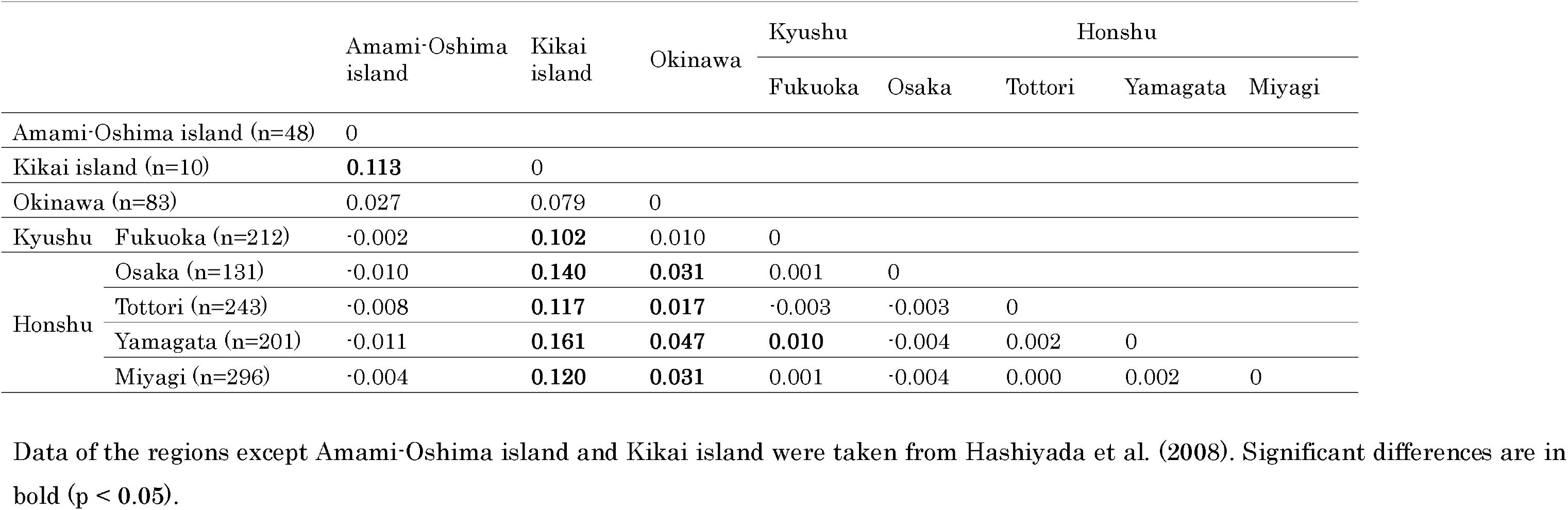
Pairwise *R*_*ST*_ values of Y-STR

### 3.3. Autosomal SNPs

Genotypes of 91 subjects whose all four grandparents were born in the Amami islands were determined, and allele frequencies were calculated.

The frequency of the T allele in rs3827760 (*EDAR*) in Amami-Oshima island (0.385) was significantly higher than in the Hondo cluster (0.222) (*P* < 0.001) and not significantly different from the Ryukyu cluster (0.398) (*P* = 0.426). The frequency of the T allele in Kikai island (0.647) was significantly higher than in Amami-Oshima island, the Ryukyu cluster, and the Hondo cluster (*P* = 0.005, *P* = 0.004, and *P* < 0.001, respectively) (Table 4).

**Table 4.**
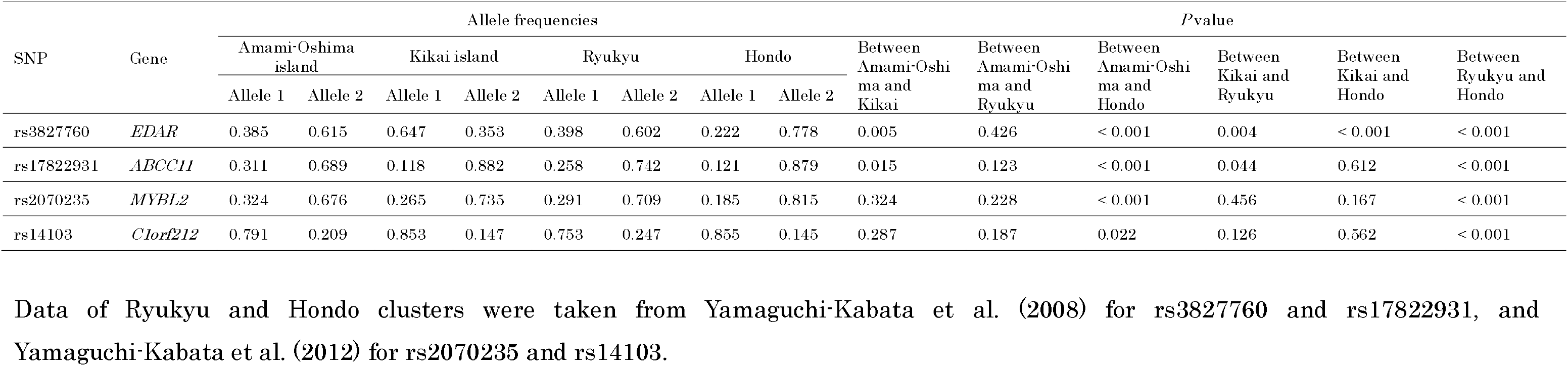
Allele frequencies of four autosamal SNPs

The frequency of the G allele in rs17822931 (*ABCC11*) in Amami-Oshima island (0.311) was significantly higher than in the Hondo cluster (0.121) (*P* < 0.001) but not significantly different from that in the Ryukyu cluster (0.258) (*P* = 0.123). The frequency of the G allele in Kikai island (0.118) was significantly lower than in the Ryukyu cluster (*P* = 0.044) and Amami-Oshima island (*P* = 0.015) but not significantly different from that in the Hondo cluster (*P* = 0.612) (Table 4).

The frequency of the G allele in rs2070235 (*MYBL2*) in Amami-Oshima island (0.324) was significantly higher than in the Hondo cluster (0.185) (*P* < 0.001) but not significantly different from that in the Ryukyu cluster (0.291) (*P* = 0.228). The frequency of the G allele in Kikai island was not significantly different from those in Amami-Oshima island, the Ryukyu cluster, and the Hondo cluster (*P* = 0.324, *P* = 0.456, and *P* = 0.167, respectively) (Table 4).

The frequency of the A allele in rs14103 (*C1orf212*) in Amami-Oshima island (0.791) was significantly lower than in the Hondo cluster (0.855) (*P* = 0.022) and not significantly different from that in the Ryukyu cluster (0.753) (*P* = 0.187), whereas that in Kikai island was not different from those in Amami-Oshima island, the Ryukyu cluster, and the Hondo cluster (*P* = 0.287, *P* = 0.126, and *P* = 0.562, respectively) (Table 4).

## 4. Discussion

Amami islanders showed a trend of Tajima’s *D* values of mtDNA to be negative. This trend is commonly observed in the populations of East Asia, including main-island Okinawa, Miyako island, Ishigaki island, and Honshu (Oota et al., 2002; Matsukusa et al., 2010). Purifying selection or recent population expansion would generally be a reason for the negative Tajima’s *D* values (Tajima, 1989), but the former is unsuitable in this case because D-loop is a non-coding region. Therefore, this result suggests the past population expansion and/or gene flow from the surrounding regions, *i.e.,* the Amani islanders were genetically not isolated for long.

Pairwise *F*_*ST*_ values and phylogenetic networks represent close genetic relations in the female lineages between Amami-Oshima and Kikai islands. The female lineages in Amami-Oshima island differed from Honshu, main-island Okinawa, and Ishigaki island, and those in Kikai island differed from Ishigaki island. In the phylogenetic network, the female lineages in Amami-Oshima and Kikai islands were apart from those in main-island Okinawa and Ishigaki island. Conversely, the female lineages in Amami-Oshima and Kikai islands did not differ from those in Miyako island, and those three were located close to each other in the phylogenetic network; however, no historical records that link geographically separated people in the Amami islands and Miyako island have been recognized. Furthermore, female lineages of Korea were not different from those of Amami-Oshima island, Kikai island, Kyushu, and Honshu, whereas female lineages of Ainu, Taiwan Han, and Taiwan aborigines were different from any other populations; this indirectly suggests that female lineages in the Amami islanders have been genetically affected by mainland Japanese strongly compared with the genetic effect of mainland Japanese on female lineages of Okinawa.

Gene diversities and *R*_*ST*_ values of the four Y-STRs indicated that the male lineages in Amami-Oshima island were located between those in Honshu and Okinawa, but those in Kikai island showed some genetic deviation from these three populations, including Amami-Oshima islanders; this is assumed to be due to the founder effect that refers to the phenomenon that gene frequencies of a small population migrated to a new location sometimes differ from those of the original population because of chance bias (Mayr, 1954). In Kikai islanders, the small population size (Kagoshima Prefecture, 2018b) and the degree of contacts with mainland Japanese different from Amami-Oshima islanders (Takanashi, 2009; Yoshinari, 2011; Takamiya, 2013; Takamiya et al., 2019) may have caused that.

Comparing the maternal (mtDNA) and paternal (Y-STR) genes (Table 5), we found that the mean of pairwise *R*_*ST*_ values of Y-STR had a trend that the larger the value is, the smaller population size is (0.034 in entire Japan, 0.055 in the Ryukyu Archipelago and Kyushu, 0.073 in the Ryukyu Archipelago, and 0.113 in the Amami islands). Conversely, the mean *F*_*ST*_ values of mtDNA did not change so much (0.020 in entire Japan, 0.021 in the Ryukyu Archipelago and Kyushu, 0.019 in the Ryukyu Archipelago, and − 0.007 in the Amami islands). These values suggest that males are genetically more divergent than females because of patrilocality; in fact, female migration was suggested frequently in the Ryukyu Archipelago (Matsukusa et al., 2010), and we confirmed this in the Amami islands.

**Table 5.**
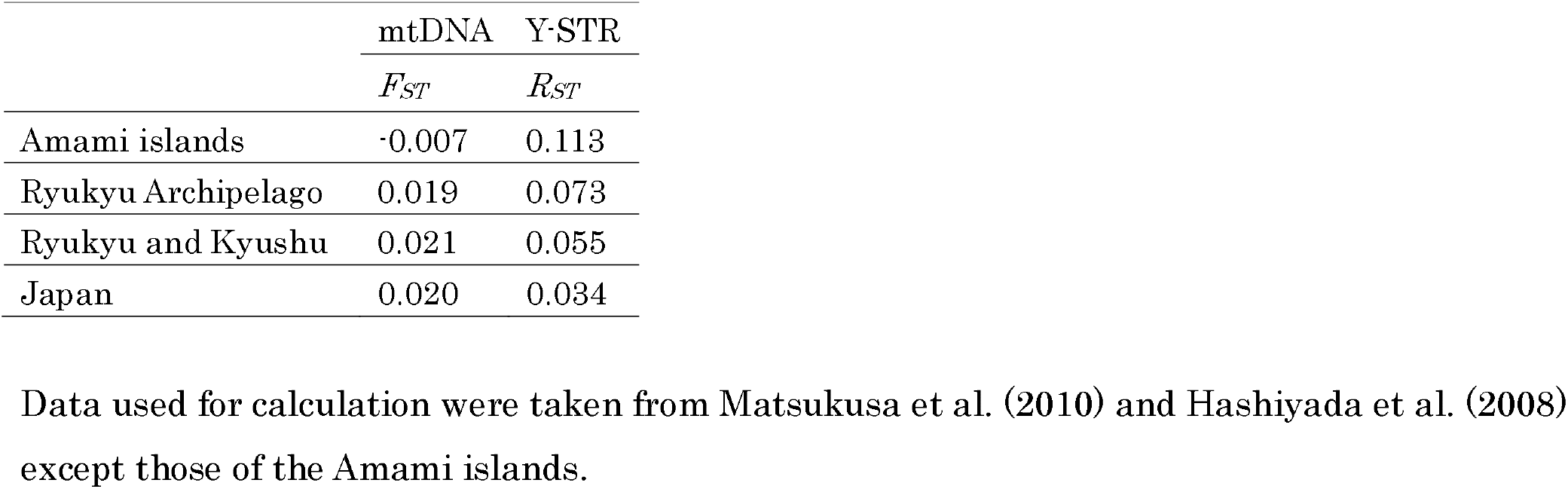
Mean of pairwise *F*_*ST*_ values of mtDNA and pairwise *R*_*ST*_ values of Y-STR

Four autosomal SNPs whose allele frequencies differed between Okinawa and the rest of Japan (Yamaguchi-Kabata et al., 2008) were selected to identify the Amami islanders’ genealogic position and search for the traits of selective pressure if it ever exerted in these populations. The allele frequencies of the four SNPs in Amami-Oshima islanders were similar to the Ryukyu cluster and significantly different from the Hondo cluster, which may be because of gene flow from Okinawa. Conversely, in Kikai islanders, the T allele frequency of rs3827760 (*EDAR*) was significantly higher than those in the other three populations, and the G allele frequency of rs17822931 (*ABCC11*) was significantly lower than those in the Ryukyu cluster and Amami-Oshima islanders but not lower than that in the Hondo cluster. These can be explained by the founder effect, as observed in Y-STR: DYS438. The presence of selective pressure is hard to recognize in these loci for the following reasons.

The C allele of rs3827760 on the EDAR gene is associated with thicker hair (Fujimoto et al., 2008a, b), straighter hair (Tan et al., 2013), multiple dental traits (Kimura et al., 2009; Park et al., 2012; Kamberov et al., 2013), and high density of sweat gland (Kamberov et al., 2013). The frequencies of the C allele are very high in East Asians and Native Americans, whereas that allele is almost absent in Europeans and Africans, and this implies strong positive selection in Asia (Sabeti et al., 2007; Bryk et al., 2008; Xue et al., 2009; Grossman et al., 2010). Adaptation to high humidity is assumed to be selective pressure (Kamberov et al., 2013; Tan et al., 2013), but frequencies of the C allele in the Ryukyu cluster, Amami-Oshima island, and Kikai island were significantly lower than that in the Hondo cluster, despite hot and humid climate in the Ryukyu Archipelago (Table 4). This suggests that selective pressure on the C allele has not been so strong in the Ryukyu Archipelago, and low frequencies of the C allele in the Amami islands may be due to gene flow from Okinawa.

Secondly, G/G and G/A genotypes of rs17822931 located on the ABCC11 gene are associated with wet earwax (Yoshiura et al., 2006) and axillary osmidrosis (Nakano et al., 2009), which are assumed to be due to their enhanced function of the apocrine gland. The frequencies of the A allele are very high in East Asia. They decline when the latitude is getting lower in Asian, African, and European populations, and this implies strong positive selection and can be explained by adaptation to cold climate, such as less sweating (Yoshiura et al., 2006; Ohashi et al., 2011). Therefore, the frequencies of the A allele are significantly lower in the Ryukyu cluster and Amami-Oshima island than in the Hondo cluster (Table 4) because selective pressure should not have existed under hotter circumstances in the Ryukyu Archipelago.

In conclusion, Amami-Oshima and Kikai island populations showed a genetically intermediate position between mainland Japan and Okinawa in mtDNA and Y-STR. However, the frequencies of several autosomal SNPs in the Amami islanders indicated a significant difference from mainland Japanese. The *EDAR* and *ABCC11* of mainland Japanese are assumed to have been under natural selection; however, those of the Amami islanders may have been affected by the gene flow from Okinawa but not natural selection. Furthermore, the extremely high or low frequencies of several alleles implied a founder effect in Kikai island. It should be noted that we could not say with strong evidence whether mtDNA and Y-STRs of the Amami islanders were close to Okinawa or mainland Japan and whether autosomal SNPs of the Amami islanders have been affected by gene flow from Okinawa because of the small sample size (especially in Kikai island) and the number of alleles in the present study. Geographically broad and detailed samplings and genome-wide analyses are awaited.

Declarations of interest: none

## Acknowledgments

We would like to thank all anonymous subjects in Naze, Amami-shi, Kagoshima, in Kasari-cho, Amami-shi, Kagoshima, in Yamato-son, Oshima-gun, Kagoshima, and in Kikai-cho, Oshima-gun, Kagoshima for participating in this study. We are grateful to Dr. M. Suzuki (Ministry of the Environment, Japan), Prof. H. Takamiya (Kagoshima University), Ms. R. Koike, and anonymous cooperators for their help in sampling. We thank Enago (https://www.enago.jp) for the English language review.

