## Supplementary material for "Genetic lineage of the Amami islanders inferred from classical genetic markers"

Supp. Table 1. Primers used for PCR and sequencing of mtDNA

| Primer name | Sequence (5'-3') | Use | Ref. |
| --- | --- | --- | --- |
| GENBANK_MT_26F | CGGCTTCGACCCTATATCC | PCR | http://insertion.stanford.edu/primers.html |
| L15996F | CTCCACCATTAGCACCCAAAGC | PCR, sequencing | Vigilant et al., 1989 |
| L16223F | AGCAAGTACAGCAATCAACC | Sequencing | Vigilant et al., 1989 |
| H16401R | TGATTTCACGGAGGATGGTG | Sequencing | Vigilant et al., 1989 |
| H408R | CTGTTAAAAGTGCATACCGCCA | PCR, sequencing | Vigilant et al., 1989 |
| GENBANK_MT_5R | CCATAGGGTCTTCTCGTCTTG | PCR | http://insertion.stanford.edu/primers.html |

Supp. Table 2. Primers used for PCR and sequencing of Y-STR

| Primer name | Sequence (5'-3') | Annealing temperature | Ref. |
| --- | --- | --- | --- |
| DYS393-F | GTGGTCTTCTACTTGTGTCAATAC | 53°C | Kayser, et al., 1997; de Kniff, et al., 1997 |
| DYS393-R | AACTCAAGTCCAAAAAATGAGG |  |  |
| DYS19-F | CTACTGAGTTTCTGTTATAGT | 53°C | Roewer and Epplen., 1992 |
| DYS19-R | ATGGCCATGTAGTGAGGACA |  |  |
| DYS19-F2 | ACTACTGAGTTTCTGTTATAGTGTTTTT | 59°C | Butler et al., 2002 |
| DYS19-R2 | GTCAATCTCTGCACCTGGAAAT |  |  |
| DYS391-F | CTATTCATTCAATCATACACCCA | 53°C | Kayser, et al., 1997; de Kniff, et al., 1997 |
| DYS391-R | GATTCTTTGTGGTGGGTCTG |  |  |
| DYS438-F | TGGGGAATAGTTGAACGGTAA | 53°C | Ayub et al., 2000 |
| DYS438-R | GTGGCAGACGCCTATAATCC |  |  |
| DYS438-F2 | CCAAAATTAGTGGGGAATAGTTG | 59°C | Butler et al., 2002 |
| DYS438-R2 | GATCACCCAGGGTCTGGAGTT |  |  |

Supp. Table 3. Primers and restriction enzymes used for PCR-RFLP of autosomal SNPs

| Primer name | Sequence (5'-3') | Annealing temperature | Restriction enzyme | Ref. |
| --- | --- | --- | --- | --- |
| EDAR-RFLP-F | AGGTCTTAGCCCCACGGAACTGCCAT | 65°C | *Fnu4H I* | Hayashida et al., 2010 |
| EDAR-RFLP-R | GGACTCCACAGCATCCAACCGCTC |  |  |  |
| ABCC11-RFLP-F | TGCAAAGAGATTCCACCAGTT | 54°C | *Dde I* | Hayashida et al., 2010 |
| ABCC11-RFLP-R | AAGGTCTTCATTTTCTAGACAGC |  |  |  |
| MYBL2-F | GGATGGCCACACCATCTCAG | 62°C | *BanI* | Present study |
| MYBL2-R | GCCAGGTCTCGTTTTGCTCA |  |  |  |
| LOC113444-F | CCAGCAGTGCACCAGTAAAC | 58°C | *Nco I* | Present study |
| LOC113444-R | GGGCCCTATGGTCCTACTGT |  |  |  |

Supp. Table 4. Haplotypes observed in the Amami islands and numbers of individuals in each region

| Ht | DYS393 | DYS19 | DYS391 | DYS438 | Amami-Oshima island (n=47) | Kikai island (n=11) | Okinawa (n=83) | Kyushu | Honshu | | | |
| --- | --- | --- | --- | --- | --- | --- | --- | --- | --- | --- | --- | --- |
|  |  |  |  |  |  |  |  | Fukuoka (n=212) | Osaka (n=131) | Tottori (n=243) | Yamagata (n=201) | Miyagi (n=296) |
| 1 | 12 | 14 | 10 | 10 | 2 | 0 | 0 | 1 | 0 | 1 | 1 | 3 |
| 2 | 12 | 14 | 10 | 11 | 1 | 1 | 0 | 4 | 6 | 5 | 6 | 7 |
| 3 | 12 | 15 | 10 | 10 | 1 | 1 | 2 | 10 | 4 | 12 | 1 | 8 |
| 4 | 12 | 16 | 10 | 10 | 5 | 0 | 2 | 8 | 3 | 9 | 7 | 8 |
| 5 | 12 | 17 | 10 | 10 | 3 | 0 | 2 | 6 | 1 | 12 | 2 | 4 |
| 6 | 13 | 13 | 10 | 10 | 1 | 2 | 1 | 0 | 0 | 7 | 2 | 3 |
| 7 | 13 | 14 | 10 | 10 | 1 | 0 | 0 | 2 | 3 | 2 | 4 | 4 |
| 8 | 13 | 15 | 10 | 10 | 1 | 0 | 3 | 13 | 12 | 10 | 16 | 28 |
| 9 | 13 | 15 | 10 | 11 | 1 | 1 | 0 | 2 | 1 | 4 | 1 | 7 |
| 10 | 13 | 15 | 10 | 13 | 8 | 0 | 10 | 38 | 21 | 31 | 43 | 51 |
| 11 | 13 | 15 | 11 | 10 | 0 | 1 | 4 | 3 | 0 | 1 | 1 | 5 |
| 12 | 13 | 16 | 10 | 10 | 2 | 0 | 10 | 17 | 4 | 10 | 13 | 24 |
| 13 | 13 | 16 | 10 | 13 | 2 | 0 | 1 | 9 | 12 | 18 | 16 | 11 |
| 14 | 13 | 16 | 10 | 14 | 2 | 0 | 0 | 0 | 0 | 0 | 1 | 1 |
| 15 | 13 | 16 | 11 | 10 | 2 | 0 | 1 | 0 | 0 | 3 | 0 | 1 |
| 16 | 13 | 17 | 10 | 10 | 1 | 2 | 16 | 24 | 14 | 23 | 25 | 32 |
| 17 | 13 | 17 | 11 | 10 | 1 | 0 | 2 | 3 | 2 | 7 | 4 | 3 |
| 18 | 14 | 13 | 10 | 10 | 3 | 0 | 6 | 6 | 4 | 8 | 6 | 2 |
| 19 | 14 | 15 | 10 | 9 | 0 | 1 | 1 | 1 | 0 | 0 | 1 | 0 |
| 20 | 14 | 15 | 10 | 10 | 3 | 0 | 0 | 5 | 5 | 12 | 5 | 8 |
| 21 | 14 | 15 | 10 | 13 | 3 | 0 | 2 | 1 | 1 | 5 | 2 | 1 |
| 22 | 14 | 16 | 10 | 9 | 1 | 1 | 3 | 4 | 0 | 0 | 0 | 1 |
| 23 | 14 | 16 | 11 | 13 | 2 | 0 | 4 | 0 | 0 | 0 | 0 | 0 |
| 24 | 14 | 17 | 10 | 9 | 1 | 0 | 2 | 1 | 0 | 0 | 0 | 0 |
| 25 | 15 | 13 | 10 | 10 | 0 | 1 | 2 | 2 | 0 | 0 | 0 | 0 |
| others |  |  |  |  | 0 | 0 | 9 | 52 | 38 | 63 | 44 | 84 |

Data of the regions except Amami-Oshima island and Kikai island were taken from Hashiyada et al. (2008).

Supp. Table 5. Allele frequencies of four Y-STRs

| DYS393 |  | | |  | | |  | | |  | | |  | | | DYS391 | | |  | | |  | | |  | | |  | |
| --- | --- | --- | --- | --- | --- | --- | --- | --- | --- | --- | --- | --- | --- | --- | --- | --- | --- | --- | --- | --- | --- | --- | --- | --- | --- | --- | --- | --- | --- |
| Alleles | Frequency | | | | | | | | | | | |  | | | Alleles | | | Frequency | | | | | | | | | | |
|  | | Amami-Oshima island (n=48) | | | Kikai island (n=10) | | | Okinawa (n=87) | | | Hounshu (n=207) | | |  | | |  | | | Amami-Oshima island (n=48) | | | Kikai island (n=10) | | | Okinawa (n=87) | | | Hounshu (n=207) |
| 11 | | 0 | | | 0 | | | 0 | | | 0.024 | | |  | | | 9 | | | 0 | | | 0 | | | 0 | | | 0.039 |
| 12 | | 0.250 | | | 0.200 | | | 0.230 | | | 0.222 | | |  | | | 10 | | | 0.875 | | | 1 | | | 0.805 | | | 0.865 |
| 13 | | 0.479 | | | 0.500 | | | 0.506 | | | 0.618 | | |  | | | 11 | | | 0.125 | | | 0 | | | 0.184 | | | 0.092 |
| 14 | | 0.271 | | | 0.200 | | | 0.218 | | | 0.126 | | |  | | | 12 | | | 0 | | | 0 | | | 0.012 | | | 0.005 |
| 15 | | 0 | | | 0.100 | | | 0.046 | | | 0.010 | | |  | | |  | | |  | | |  | | |  | | |  |
| DYS19 | |  | | |  | | |  | | |  | | |  | | | DYS438 | | |  | | |  | | |  | | |  |
| Alleles | Frequency | | | | | | | | | | | |  | | | | | Alleles | | | Frequency | | | | | | | | |
|  | Amami-Oshima island (n=48) | | | Kikai island (n=10) | | | Okinawa (n=87) | | | Hounshu (n=207) | | |  | | |  | | | Amami-Oshima island (n=48) | | | Kikai island (n=10) | | | Okinawa (n=87) | | | Hounshu (n=207) | |
| 13 | 0.083 | | | 0.300 | | | 0.081 | | | 0.048 | | |  | | | 8 | | | 0 | | | 0 | | | 0 | | | 0.005 | |
| 14 | 0.083 | | | 0.100 | | | 0.058 | | | 0.058 | | |  | | | 9 | | | 0.042 | | | 0.200 | | | 0.058 | | | 0.014 | |
| 15 | 0.375 | | | 0.300 | | | 0.299 | | | 0.488 | | |  | | | 10 | | | 0.563 | | | 0.600 | | | 0.667 | | | 0.556 | |
| 16 | 0.333 | | | 0.100 | | | 0.287 | | | 0.217 | | |  | | | 11 | | | 0.042 | | | 0.200 | | | 0.046 | | | 0.106 | |
| 17 | 0.125 | | | 0.200 | | | 0.276 | | | 0.184 | | |  | | | 12 | | | 0 | | | 0 | | | 0 | | | 0.034 | |
| 18 | 0 | | | 0 | | | 0 | | | 0.005 | | |  | | | 13 | | | 0.313 | | | 0 | | | 0.230 | | | 0.266 | |
|  |  | | |  | | |  | | |  | | |  | | | 14 | | | 0.042 | | | 0 | | | 0 | | | 0.019 | |

For the analyses of Y-STR, one subject from Kikai island whose paternal grandfather was born in Amami-Oshima island was excluded from Kikai island samples and included in Amami-Oshima island samples. Data of Honshu and Okinawa were taken from Uchihi et al. (2003).

Supp. Table 6. Gene diversities of Y-STRs

| Locus | Gene diversity | | | | *P* value | | | | | |
| --- | --- | --- | --- | --- | --- | --- | --- | --- | --- | --- |
|  | Amami-Oshima island (n=48) | Kikai island (n=10) | Okinawa (n=87) | Hounshu (n=207) | Between Amami-Oshima and Kikai | Between Amami-Oshima and Okinawa | Between Amami-Oshima and Honshu | Between Kikai and Okinawa | Between Kikai and Honshu | Between Okinawa and Honshu |
| DYS393 | 0.648 | 0.733 | 0.649 | 0.555 | 0.352 | 0.538 | 0.107 | 0.849 | 0.228 | 0.027 |
| DYS19 | 0.734 | 0.844 | 0.751 | 0.679 | 0.194 | 0.354 | 0.316 | 0.199 | 0.062 | 0.046 |
| DYS391 | 0.223 | 0.000 | 0.322 | 0.244 | 0.577 | 0.652 | 0.532 | 0.288 | 0.739 | 0.019 |
| DYS438 | 0.593 | 0.622 | 0.503 | 0.611 | 0.035 | 0.320 | 0.351 | 0.026 | 0.041 | 0.045 |

Data of Honshu and Okinawa were taken from Uchihi et al. (2003).

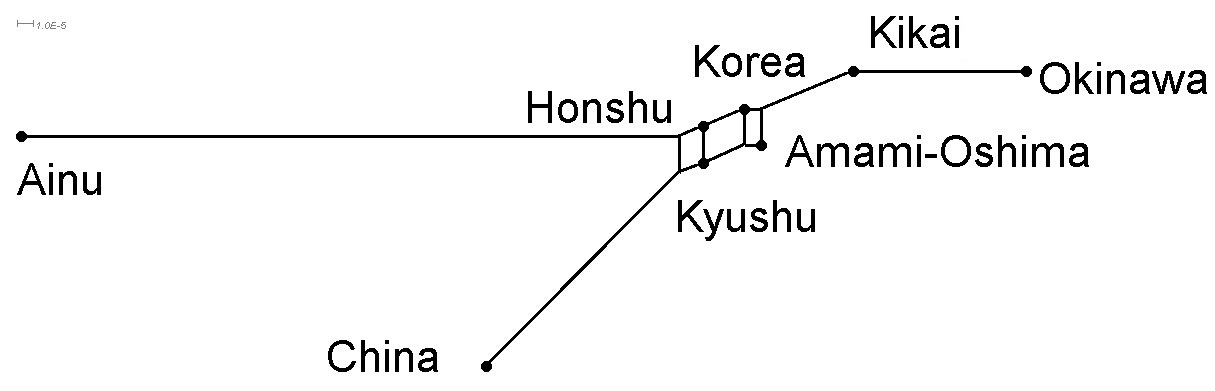

Supp. Fig. 1. Phylogenetic network between populations based on *d_A_* distances calculated from 487 bp of D-loop region. Data of the populations except the Amami islands were taken from Horai et al. (1996).
